# Temporal Generative Models for Learning Heterogeneous Group Dynamics of Ecological Momentary Data

**DOI:** 10.1101/2023.09.13.557652

**Authors:** Soohyun Kim, Young-geun Kim, Yuanjia Wang

## Abstract

One of the goals of precision psychiatry is to characterize mental disorders in an individualized manner, taking into account the underlying dynamic processes. Recent advances in mobile technologies have enabled the collection of Ecological Momentary Assessments (EMAs) that capture multiple responses in real-time at high frequency. However, EMA data is often multi-dimensional, correlated, and hierarchical. Mixed-effects models are commonly used but may require restrictive assumptions about the fixed and random effects and the correlation structure. The Recurrent Temporal Restricted Boltzmann Machine (RTRBM) is a generative neural network that can be used to model temporal data, but most existing RTRBM approaches do not account for the potential heterogeneity of group dynamics within a population based on available covariates. In this paper, we propose a new temporal generative model, the Heterogeneous-Dynamics Restricted Boltzmann Machine (HDRBM), to learn the heterogeneous group dynamics and demonstrate the effectiveness of this approach on simulated and real-world EMA data sets. We show that by incorporating covariates, HDRBM can improve accuracy and interpretability, explore the underlying drivers of the group dynamics of participants, and serve as a generative model for EMA studies.

## 1. Introduction

Mental disorders are often assessed by clinician-administered instruments that rely on a single summary score to measure disease severity. For example, the Hamilton Rating Scale for Depression (HAM-D; Hamilton, 1960) is commonly used to assess the severity of depression. However, mental illnesses are complex and multifaceted; individual symptoms differ in their associations to risk factors (Fried and Nesse, 2015) and impact on patient functioning (Tweed, 1993), and therefore are not interchangeable. Thus, instead of solely relying on a summary score, a more sophisticated and nuanced approach is to jointly model how individual symptoms arise from multiple underlying latent constructs under measurement theory and statistical methods (Lohr, 2002; Bollen, 2002). This approach takes into account that individual symptoms may not be interchangeable and aims to capture the true mental states of interest through latent models. The unobserved latent constructs reflect the true mental states of interest and can be measured through clinical instruments.

Traditionally, clinical instruments and behavioral tests for assessing mental disorders are collected at given time points in a clinical or counseling setting, which does not continuously capture patients’ moods or behaviors in real-time, and it may be influenced by recall bias (Shiffman et al., 2008). Advancements in technology, such as mobile devices, have enabled the collection of comprehensive and real-time data on patients’ moods and behaviors to overcome some of the limitations of traditional mental health assessments. Ecological Momentary Assessment (EMA) is a collection of assessments of subjects’ current or recent mental states, repeatedly measured over time, in their natural environments (Shiffman et al., 2008). EMA can provide more accurate and comprehensive observations of patients’ symptoms and functioning, capturing their experiences and behaviors as they occur in real life. This can help identify patterns and triggers of symptoms or emotions and track changes in response to interventions over time.

Nonetheless, the analysis of EMA data is challenging. First, patients affected by mental disorders present substantial heterogeneity in their symptomatology. For example, major depressive disorder (MDD) can be diagnosed based on 227 different symptom combinations (Fried and Nesse, 2015), and the course of the disease can vary significantly between patients (Wakefield and Schmitz, 2014). Methods that are flexible to account for extensive heterogeneity and individual-level variation are desirable. EMA data can be regarded as intensively measured longitudinal data with a large number of measures and time points. As in longitudinal data analysis, both between-subject variation and correlation between observations within-subject need to be addressed. In addition, different subjects’ EMA experiences can vary widely, and the measurement patterns can be highly unbalanced. Therefore, methods that can handle complex temporal dependence and potentially long-range correlation are needed.

A common method for analyzing EMA data is data aggregation, for example, summarizing an item in EMA data over time into a single value such as the mean or variability (e.g., the Root Mean Square of Successive Differences; Shaffer and Ginsberg (2017)). However, these methods may overlook the detailed patterns regarding the evolution of a patient’s time-course experience. Alternative methods beyond simple aggregation are based on the hierarchical linear model, also known as a multilevel model or linear mixed-effects model. In usual mixed-effects models, both within-subjects (WS) variance and between-subjects (BS) variance are assumed to be homogeneous across subjects, which does not address heterogeneity observed in many EMA studies. An extension is the location-scale mixed-effects model (Hedeker et al., 2008), where log-linear (sub)models are included for both WS and BS variance, allowing covariates to influence both sources of variation. The model also includes a random subject-specific effect on the WS variance specification. This permits the WS variance to vary at the subject level beyond the influence of covariates on this variance. The most recent extension of mixed-effect models developed for analyzing EMA data includes the latent multivariate location-scale linear mixed-effects model, i.e., location-scale LMM (Williams et al., 2021). This method can handle multiple correlated outcomes and includes multiple correlated latent variables as random effects.

The aforementioned location-scale LMMs are developed for multivariate continuous outcomes, which poses a limitation in the analysis of EMA data where the measures can be discrete. Additionally, they require parametric assumptions on the fixed and random effects as well as on the correlation structure over time, which may not be fully satisfied by EMA data. On the other hand, generalized mixed-effects models can handle discrete outcomes but often suffer from computational challenges with intensive EMA data. Machine learning methods (e.g., random forest, recurrent neural networks [RNN]) can handle complex sequence data with greater flexibility and accommodate both discrete and continuous measures. For example, Barrigón et al. (2017) used a random forest to predict active users of EMA, and Hulme et al. (2020) developed a continuous-time Hidden Markov Model where unobserved states are inferred from observed response sequences and are used to predict the probability of transition at any time in the future. Restricted Boltzmann machine (RBM; Hinton and Salakhutdinov, 2006), recurrent temporal restricted Boltzmann machine (RTRBM; Sutskever et al., 2008), and Long Short-Term Memory (LSTM) are other types of neural networks that can be used in EMA analysis (Mikus et al., 2018; Chen et al., 2021; Koppe et al., 2019). LSTM can handle complex temporal patterns, including long-term dependencies and irregular sampling intervals, and are capable of predicting both continuous and discrete outcomes. However, LSTM has limited interpretability, making it difficult to examine the roles of latent units on the observed measures. RBM and its temporal variation, RTRBM, overcome this problem by providing a clear and interpretable structure to the model using conditionally independent latent variables. This conditional independence implies that the latent variables capture distinct, non-redundant features of the data, allowing for a more meaningful interpretation of the learned representations. Yet, current variations of the RBM do not handle covariates, leaving heterogeneity unaccounted for.

In this work, to jointly model multiple sequences of categorical or ordinal measures in EMA studies as arising from lower-dimensional latent states while accounting for heterogeneity, we propose a new temporal generative model, the Heterogeneous-Dynamics Restricted Boltzmann Machine (HDRBM). Our method aligns with measurement theory and extends the RBM and RTRBM to model EMA data’s complex dependencies and temporal structure given covariates. We model the higher-dimensional observed symptoms using low-dimensional latent states that can capture the most important variations of patients’ underlying mental states in a multi-domain space. HDRBM is more flexible than generalized mixed-effects models and avoids their computational instability for intensively-measured EMA data. Compared to HMM, which requires Markov assumptions, our method allows the hidden state probabilities from previous time points to impact the future and thus accounts for long-range time dependence. Compared to LSTM, our method is more interpretable in that conditionally independent hidden nodes are meaningful and that the generation process is identifiable, i.e., the learning distribution of observed symptoms implies recovering the underlying latent states. Lastly, compared to RTRBM, HDRBM is able to account for the heterogeneous dynamics of patients’ EMA data. To demonstrate these advantages, we conduct simulation studies to show that HDRBM can recover the underlying latent structures and adequately reconstruct EMA data. Finally, we apply the HDRBM to a real-world EMA study to examine whether patients’ EMA experiences are associated with their mental health diagnoses at baseline. Results demonstrate that our method captures heterogeneous dynamics that differ by diagnostic groups and that the performance is much improved over mixed-effects model-based methods.

The remainder of this work is organized as follows. In Section 2, we review RBM, RTRBM and propose a new temporal generative model with covariates, HDRBM, to learn heterogeneous group dynamics. In Section 3 and Section 4, we present numeric studies through simulations and analysis of real-world EMA data, respectively. Through these studies, we demonstrate two notable properties of the proposed HDRBM: (i) the numeric stability of parameters; and (ii) the ability to capture heterogeneous temporal dynamics by diagnostic groups in the EMA data. Finally, we summarize our main contributions and future directions in Section 5.

## 2. Methodology

### 2.1 RBM for Multivariate Categorical Measures

We first provide a review of RBM and RTRBM. Let 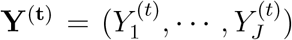 represent the *J* observable or visible units (e.g., severity rating of each symptom in HAM-D or a discrete measure in the EMA data) and 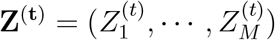 the *M* binary latent (i.e., hidden) units at time *t ∈ {*1, …, *T }* where *T* is the total number of time points (we will use the terms “latent” and “hidden” interchangeably henceforward). Each visible unit at time *t*, 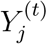, is a categorical or ordinal variable with *P* levels, such that 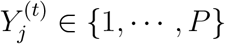, for all *j* = 1, …, *N* . The original RBM was proposed for binary visible units (Hinton and Salakhutdinov, 2006). RBM can also model categorical variables as in Salakhutdinov et al. (2007), which used softmax visible units for collaborative filtering of recommender systems. 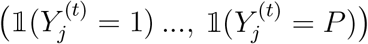, be a “one-hot” encoded vector of 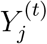, where *j* = 1, …, *N* and *t* = 1, …, *T* . For convenience of notation, we will use 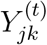 to represent 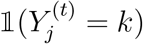 henceforward. Other more parsimonious parametrization that considers the ordinal structure is possible under the proportional odds assumption but comes at the price of a stronger assumption that may not be supported by empirical data (e.g., Section 4). We present our method under the more general multinomial model framework, in which the effects of a hidden node activation on the odds of a visible node may vary across the different levels of the visible node.

The RBM is a generative model that can learn a joint probability distribution over high-dimensional discrete observed variables and a set of latent variables, which are typically binary. The RBM has two layers: an input layer consisting of visible nodes, which correspond to the observed variables, and a hidden layer consisting of hidden units, which correspond to the latent variables (See Web Figure 1). Specifically, considering observations at a single time point, the RBM is a two-layer undirected graphical model that represents a joint distribution over visible units, **Y**, and binary hidden units **Z**, taking the form:

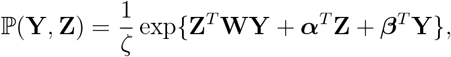

where **W** are the weights, interaction parameters between each visible and hidden unit, ***α*** and ***β*** are hidden and visible bias parameters, respectively, and *ζ* =Σ_**Y**_Σ_**Z**_exp*{***Z**^*T*^**WY**+***α***^*T*^ **Z** + ***β***^*T*^ **Y***}* is the normalization constant.

The RBM framework has connections only between the layer of hidden and the layer of visible variables but not between any two variables of the same layer. This implies that the observed variables are conditionally independent, given the state of the hidden variables, and vice versa. This characteristic is consistent with assumptions in many measurement models (e.g., in factor analysis, observed variables are independent given latent constructs). Conditional independence allows efficient learning by simplifying the computation of probabilities and parallel processing of hidden nodes as they can be updated independently. The structure of an RBM allows it to learn efficient representations of the data, making it useful for tasks such as dimensionality reduction and collaborative filtering. The basic underlying idea is that hidden units of a trained RBM represent relevant features of the observations, and these features can be used as input for other RBMs.

When applied to EMA studies of mental disorders, hidden units in the RBM are analogous to latent variables representing patients’ underlying latent mental status, such as positive or negative moods. Since the latent states **Z**^(**t**)^ are unobserved, they are learned from training on the input data **Y**^(**t**)^, which are the multiple-domain symptom items collected in EMA studies. The parameters such as **W** reflect interactions between hidden and visible units, which are analogous to factor loadings in the measurement models. Due to the measurement invariance principle in psychometrics (Meredith, 1993), the loadings (e.g., **W**) are intrinsic to an instrument and do not change over time or context, that is, an instrument measures the same construct, but the latent constructs, or hidden units in our case, may vary across groups and time.

### 2.2 Accounting for Temporal Structure and Covariates

The RBM is the building block of RTRBM (Sutskever et al., 2008) and our proposed model, HDRBM. RTRBM is a temporal generative neural network model consisting of stacked RBMs, in which the state of one or more of the previous time points modulates the states of hidden units at the subsequent time points. Each set of **Y**^(**t**)^ and **Z**^(**t**)^ at fixed time *t* is itself an RBM structure; in particular, it is a conditional RBM which depends on hidden state probabilities obtained from a recurrent neural network (RNN) (Mittelman et al., 2014). To handle temporal structure, RTRBM has an additional component to model two consecutive hidden states as a function of the visible and hidden states at previous time points and can modulate the latent states of future time points.

Building on RTRBM, we propose HDRBM to use additional covariates to model heterogeneous group dynamics over time. The incorporation of covariate information can improve the interpretability and generalizability of the model, as well as its predictive accuracy. For example, in mental health research, covariates such as demographic information, medication use, and clinical diagnosis can be used to examine the drivers that influence patients’ mental health profiles and to identify subgroups of patients with different treatment needs.

Specifically, let *X* denote any baseline covariates that may affect how latent states change over time. There is no technical difficulty in extending our method to time-varying covariates, but it will be omitted to simplify the presentation. As illustrated in Figure 1, covariate *X* modulates the latent state probabilities at *t* ⩾ 2, and its interaction effect on hidden states over time is modeled through the additional parameter *V*. The associated joint distribution of visible and hidden units given covariate *X* over all time points is

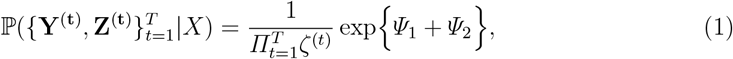

where

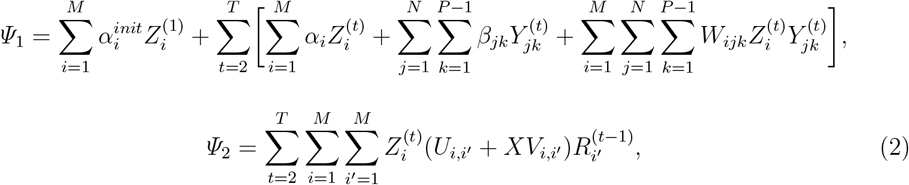

and 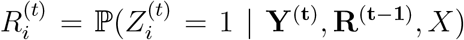, i.e., the conditional probability of hidden node *i* activation at time *t* and 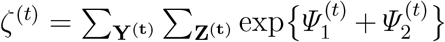 is the normalization constant at time *t* (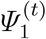 and 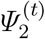 are defined in Section 2.3).

**Figure 1:**
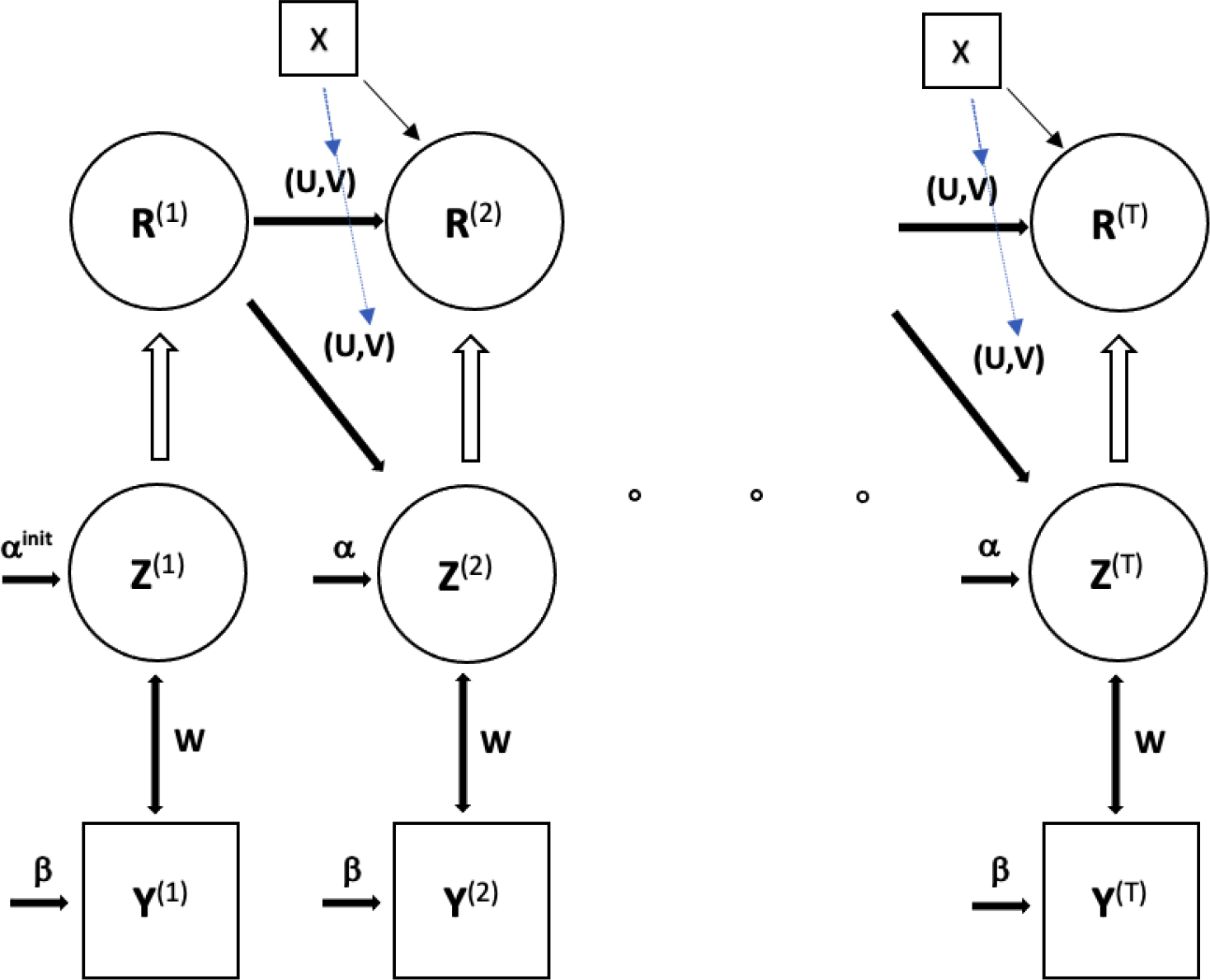
Graphical representation of the HDRBM

The parameters **W, *α*** and ***β*** are the weights and biases that are shared across all time points, *t* = 1, …, *T* . The HDRBM network now also has the following additional parameters: **U, V**, and ***α***^***init***^·**U** are the weights associated with any two consecutive hidden states for the baseline group or the average population (e.g., *X* = 0 for a binary covariate or a centered continuous covariate) and thus can be viewed as a temporal interaction parameter, **V** captures temporal heterogeneity between consecutive hidden states associated with a covariate, and ***α***^***init***^ are the initial hidden biases. In the HDRBM framework, hidden variables at time *t* are conditionally independent given the visible units at time *t* and hidden unit probabilities at previous time points. Specifically, the corresponding visible and hidden conditional distributions for the HDRBM model are, respectively,

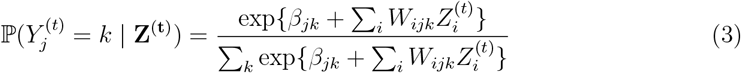

and

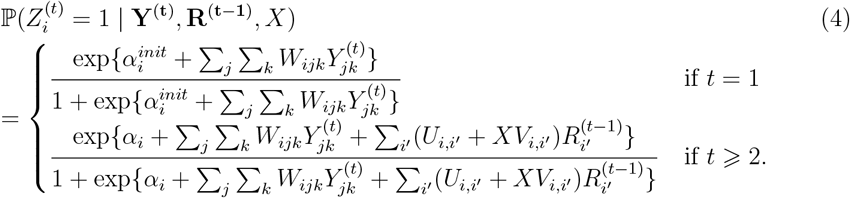

Note that the above models satisfy the measurement invariance principle, i.e., how an instrument measures latent mental states remains stable and does not vary over time or groups (Meredith, 1993; Bollen, 2002). Thus, model (3) describes the distribution of the item response at time *t* given hidden units as independent of the previous time point’s hidden probability, and the parameters governing this item response probability do not change over time. In addition, the dynamics of the observed items (i.e., visible units) are manifested through the time-evolving latent variables (i.e., hidden units). The conditional distribution of hidden units given visible units, as described in (4), models the time-varying nature of the mental states, and the additional parameters in **V** capture the between-group heterogeneity. The introduction of covariates in *Ψ*_2_ in (2) accounts for the heterogeneity of the dynamic item response distributions for subjects with different covariates. Specifically, note that the model (2) under the HDRBM allows the temporal dependence between previous time hidden units and the subsequent hidden units to vary between subjects with different covariates and thus induces long-range dependencies that can differ between subjects. Such flexibility will effectively account for heterogeneous between-group time-varying patterns observed in the motivating study introduced in Section 4. The model will learn how covariates (e.g., diagnosis groups) modify the dynamics from previous time points to the activation of latent states at subsequent time points. The weights (e.g., **W** in Figure 1) between latent units and observed units are assumed to be homogeneous under the invariance principle (Meredith, 1993; Chen et al., 2021). Thus, the parameters at the initial time point are also homogeneous across groups. In applications where further modeling of heterogeneity at the baseline is of interest, additional terms involving covariates can be introduced at the cost of estimating more parameters.

The latent states **Z**^(**t**)^ can be regarded as a unique representation of the latent mental states of patients, and the symptom items are measures or manifestations of latent states. Using **Z**^(**t**)^, we can cluster heterogeneous presentations of patient symptoms into homogeneous subgroups while capturing correlations among measures in **Y**^(**t**)^. Different profiles of **Z**^(**t**)^ may inform patient stratification and personalized treatment planning. For example, to examine whether patients who have similar symptom patterns may benefit from similar treatment.

### 2.3 Implementation

We provide details on the implementation of HDRBM. The usual approach to estimate the parameters in a posited model is through maximum likelihood. Under the HDRBM, where ***Θ*** is the set of all parameters in the model, the observed data likelihood for a single patient is given by:

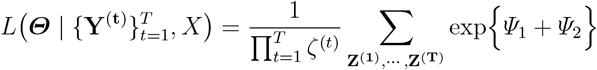

where 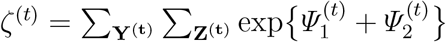 is the normalization constant for RBM at *t*, and

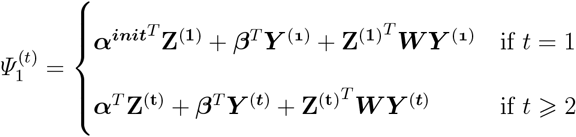

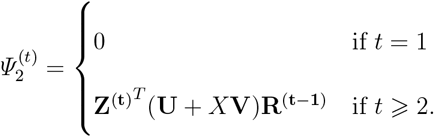

We adapt gradient-based contrastive divergence learning (Hinton, 2002), an effective tech-nique to approximate the score, or the gradient of the log-likelihoods, derived from energy densities (Qiu et al., 2019). The score of energy densities is the difference between two expected gradients of energies, called *contrastive divergence* (CD), (i) one is based on the model joint distribution of hidden and visible units, and (ii) the other is based on the product of the marginal distribution of observed visible units and the model distribution of hidden given visible units. The CD learning samples data from (i) and (ii) with Gibbs sampling to approximate the score.

Within the HDRBM context, we extend energy densities in RTRBM to (1) to reflect the heterogeneity with respect to covariates, consequently (i) and (ii) are modified into conditional ones. Specifically, 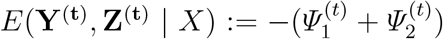is the energy and the score can be expressed as

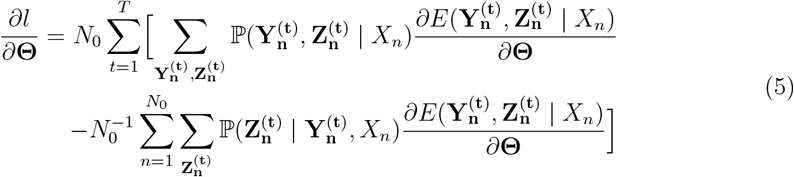

which is the difference between two expected gradients of conditional energies up to the constant multiplication. Here, *l* denotes the log-likelihood of HDRBM for *N*_0_ subjects, and *∂l/∂***Θ** denotes corresponding scores. In (5), the first term is the conditional extension of (i), based on the model joint distribution of (**Y**^(**t**)^, **Z**^(**t**)^) given *X*, and the second term is the conditional extension of (ii), based on the product of the empirical distribution of **Y**^(**t**)^ given *X* and the model distribution of **Z**^(**t**)^ given (**Y**^(**t**)^, *X*). The first term can be estimated samples obtained from a *k*-step CD, as in line 11 of Algorithm 1; the second term is estimated using empirical data. Compared with the CD learning for RTRBM, both gradients of energies and distributions are modified to conditional ones, and we sample data from two conditional distributions with Gibbs sampling to address these modifications. We use one-step Gibbs sampling, as this has previously been shown to be sufficient (Hinton, 2002). Details of the CD learning for training HDRBM are provided in Algorithm 1.

#### Algorithm 1 *k*-Step Contrastive Divergence Training for HDRBM (Mini-Batch Optimization with Momentum SGD)

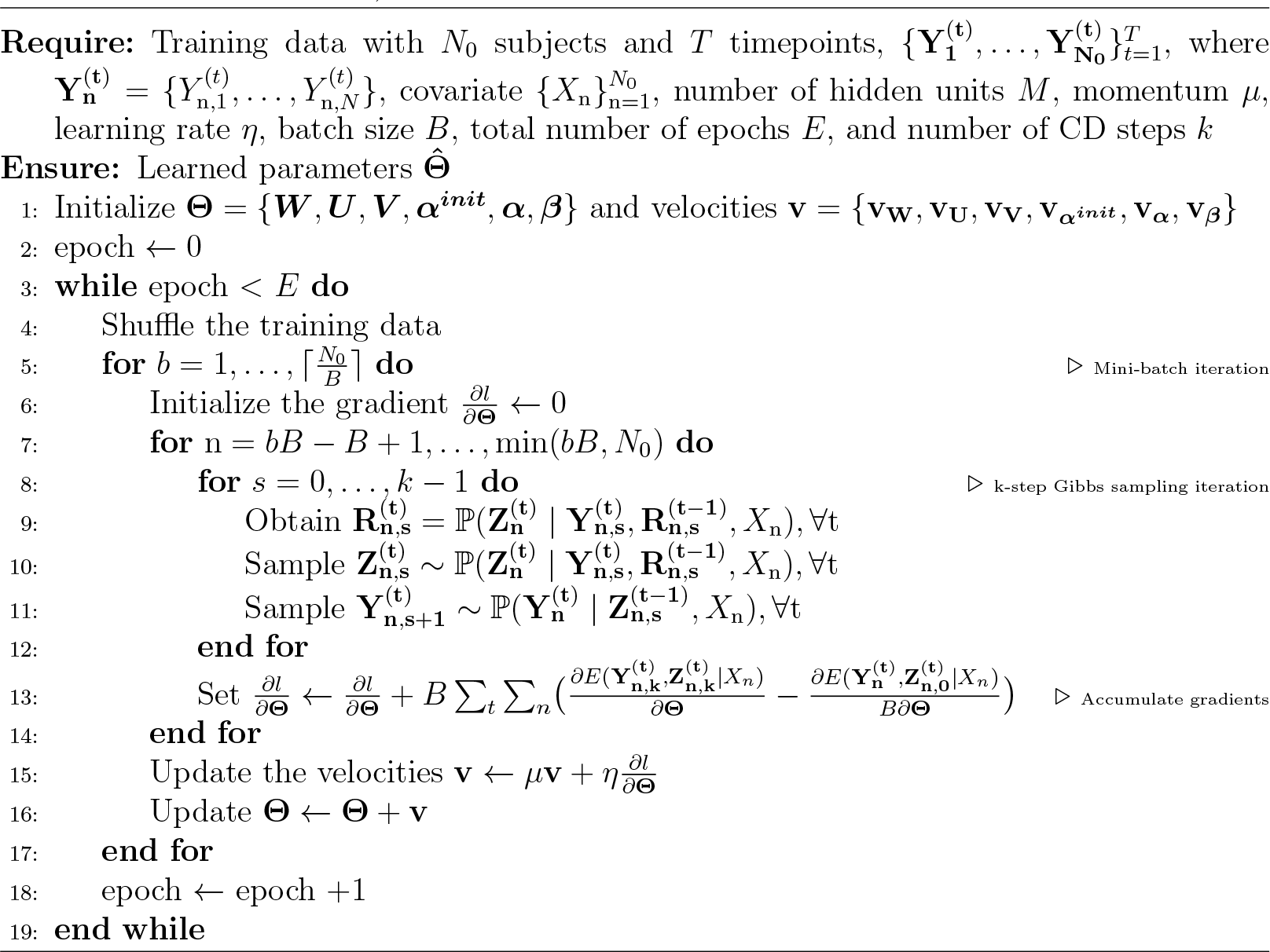

## 3. Simulation Studies

We conducted simulation studies to investigate the performance of HDRBM in terms of reconstruction accuracy and ability to recover the ground truth parameters. To generate simulated data that closely represent real world applications, we first estimated parameters of HDRBM from the real-world EMA data (Section 4). Next, using these parameters as ground truth, we applied Gibbs sampling to generate simulation datasets with varying sample sizes. We also generated a test data set of sample size 1, 000 under the same distribution. The ground truth values of parameters were denoted as 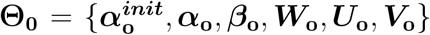 (see Section 4 and Web Tables 1-4). The simulation datasets were generated to have a known joint distribution, as specified by equation (1), with 7 time points (days) and 25 measures. We simulated four different sample size settings: *N* = 50, 100, 200, and 500. We generated and trained 100 simulation datasets in each sample size setting until stabilization of reconstruction accuracy, which took about 100 epochs, with a learning rate, weight decay, and momentum coefficient of 0.1, 0, and 0.5, respectively. Batch size varied by sample size setting; batch sizes of 5, 10, 20, and 50 were used for sample size settings of 50, 100, 200, and 500, respectively.

We found that the parameters estimated from HDRBM are close to their true values; for example, see true parameters ***W***_***0***_ and estimated weights ***Ŵ*** in Web Figure 3 and 4. Furthermore, in Figure 2, we have shown that the parameter estimates’ mean squared errors (MSEs) decrease with increasing sample size. Mean training and test accuracies of HDRBM across the 100 simulated datasets by sample size are shown in Web Table 5. These results suggest that training and testing accuracy are around 57% across all sample size settings, which is also close to the Bayes error, also as shown in Web Table 5. In conclusion, the identifiability of HDRBM allows us to recover the true underlying latent structures.

**Figure 2:**
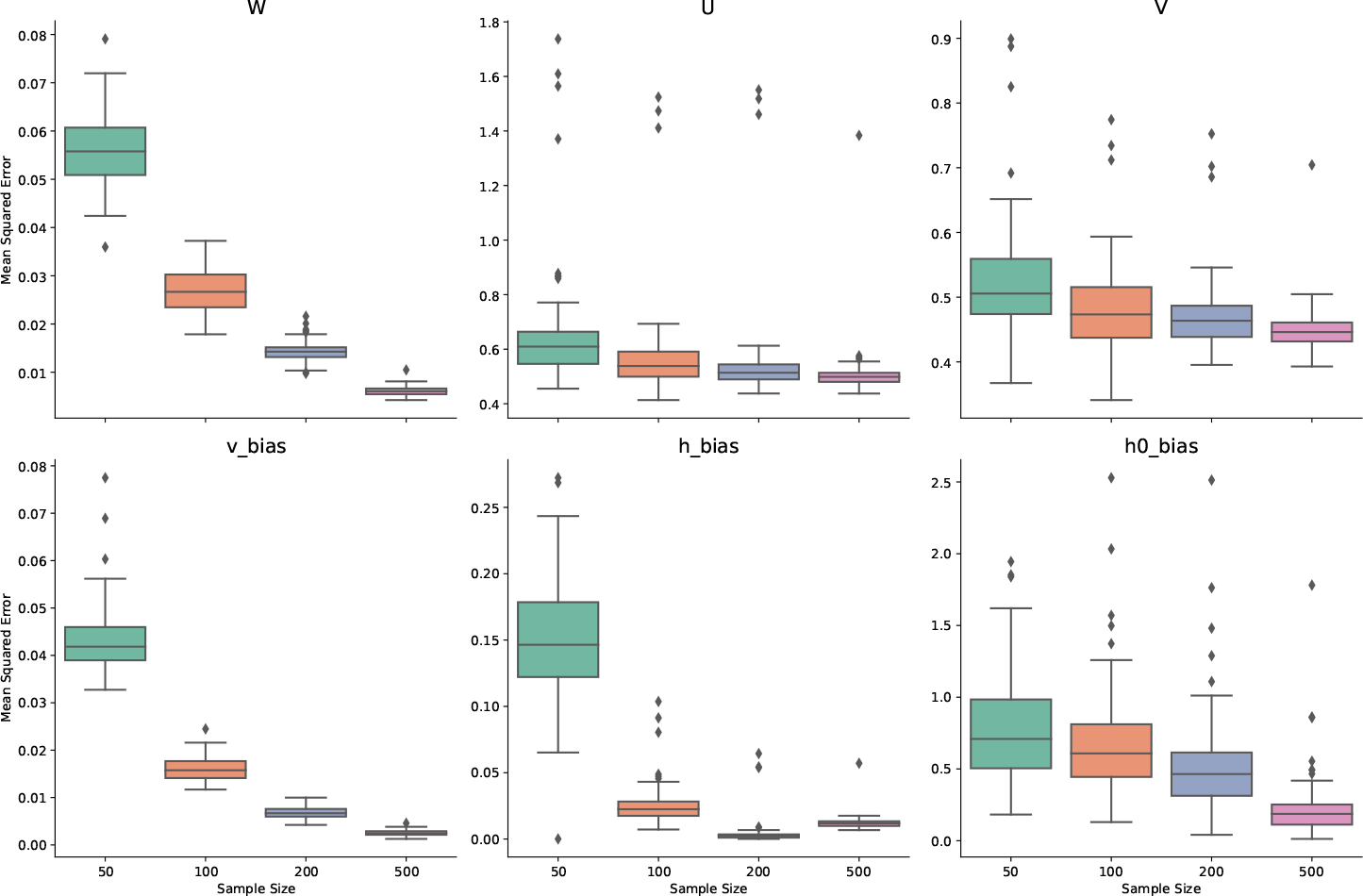
MSE of parameters estimated from HDRBM under various sample sizes (Note: outliers for **W** have been excluded for better visualization).

**Figure 3:**
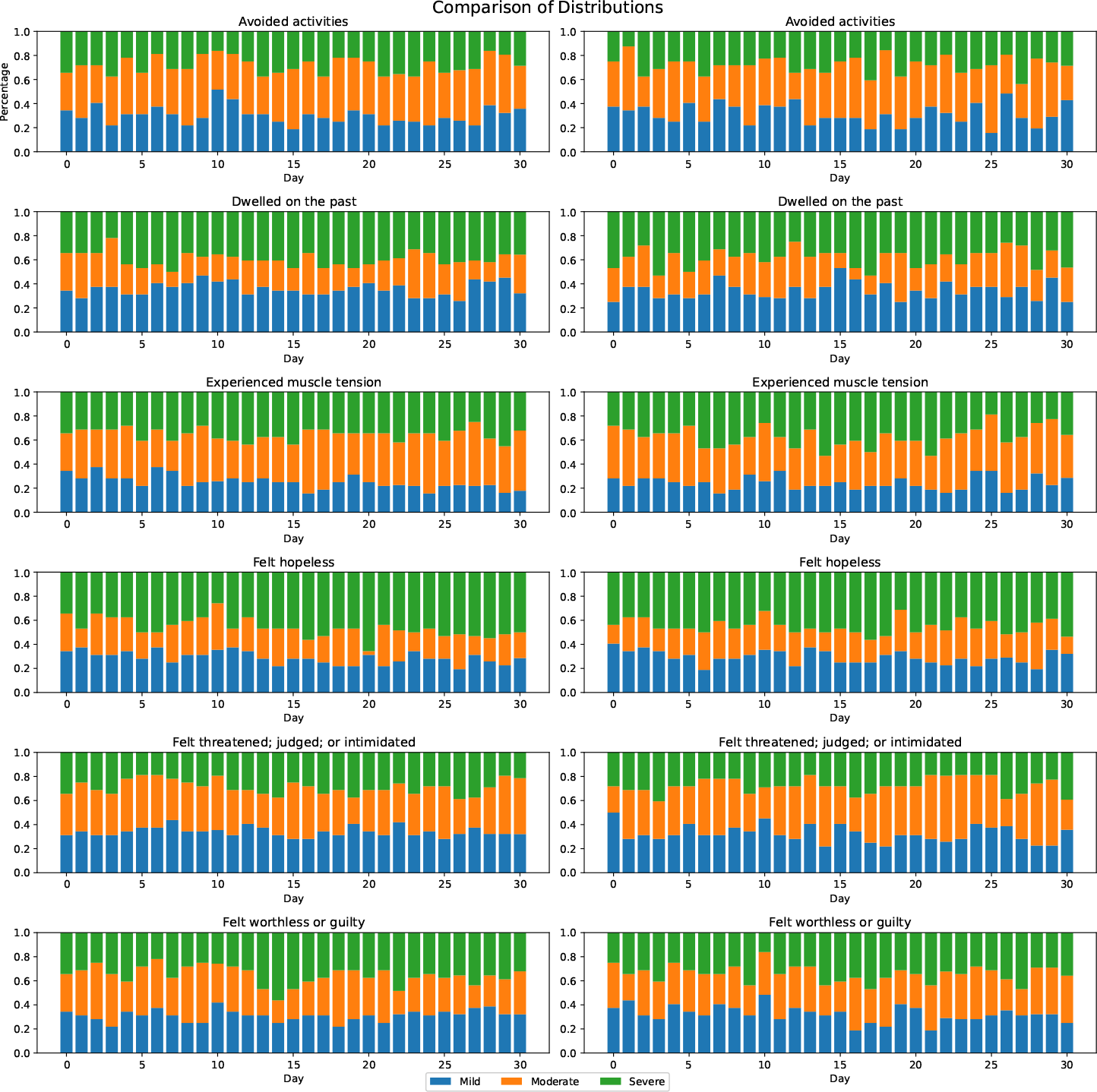
Comparison of the distribution of observed data and reconstructed data (Left: observed data; Right: HDRBM-generated data).

**Figure 4:**
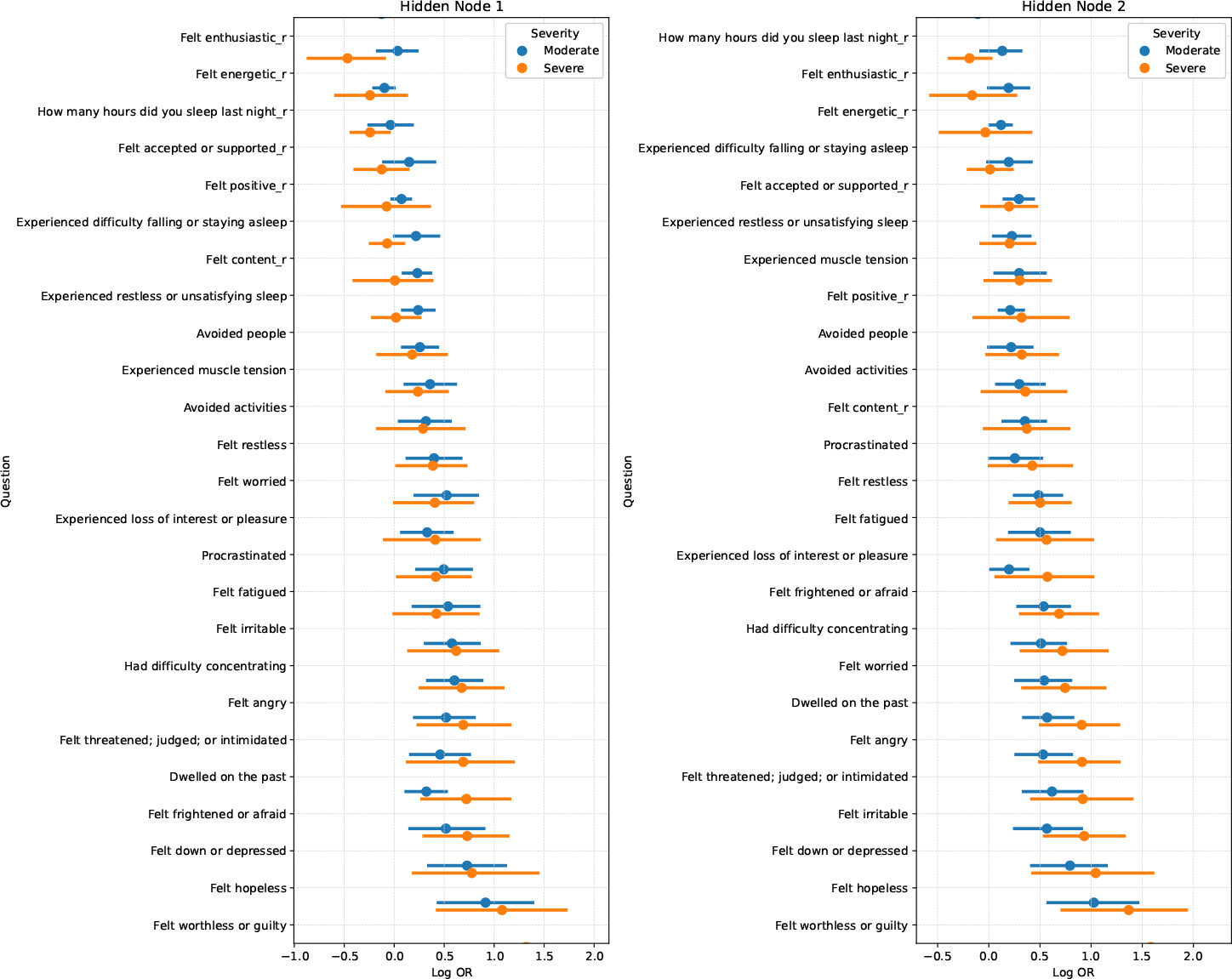
95% bootstrap confidence intervals of the log odds ratios (reference level is ‘Mild’) in the analysis of EMA data.

## 4. Analysis of the EMA data

### 4.1 Motivating study

The motivating study of this work is an EMA study conducted to identify discrete mood profiles from time-series EMA data and predict mood from such profiles (Fisher and Bosley, 2020). EMA data are becoming more prevalent as technology has enabled clinical psychologists to collect real-time self-reports of mood and behavior in subjects’ natural setting over time rather than data in a laboratory setting, which is seldom a true reflection of behavior and mood (Shiffman et al., 2008). This is important because mood is a complex construct that can be difficult to assess accurately using traditional assessments. By collecting real-time data, researchers can better understand how mood fluctuates over time and how different contextual factors and diagnostic groups influence it. EMA data are increasingly used in clinical research and practice to monitor symptoms, inform treatment, and develop personalized interventions.

The study (Bosley et al., 2019) consisted of 32 participants with primary diagnoses of a generalized anxiety disorder (GAD, *n* = 19), major depressive disorder (MDD, *n* = 6), or both (COM, *n* = 18) who were enrolled in an open trial of a personalized cognitive-behavioral intervention for mood and anxiety disorders. Individuals qualified to participate in the study and diagnosed by completing a structured clinical interview provided intensive repeated-measures data via EMA for a month before receiving psychotherapy (Fisher and Bosley, 2020; Bosley et al., 2019).

The EMA data were collected four times a day, approximately every four hours, for at least 30 days. For our analysis, we included participants’ data on the first 30 days, where for each participant, the first EMA data submitted was considered the baseline values (Day 0). 25 questions related to mood and experience were collected for each survey, for which participants submitted a score of range 0 (not at all) to 100 (as much as possible) for each question (Fisher 2019). Data for each patient were averaged over four pings submitted daily to calculate a single score for one day per patient. We created discrete EMA measures to be consistent with the common measures used to assess depression (e.g., HAM-D) and positive and negative emotion (PNASS; Wright et al., 2019) by discretizing data into 0 (mild), 1 (moderate), and 2 (severe) based on baseline tertiles. This pre-processing step improves accuracy, robustness, and reproducibility due to the noise in the EMA ratings and offers opportunities to compare with existing literature analyzing depression symptoms. The final analysis dataset included 24,800 observations (i.e., 992 pings) from 25 questions collected from 32 participants over 30 days.

We present distributions of severity levels of seven items between co-morbid patients (COM) with both MDD and GAD and patients with a single mental disorder, either MDD or GAD. Web Figure 2 illustrates the probability distribution of severity categories over one month for seven items, “Avoided activities”, “Dwelled on the past”, “Experienced muscle tension”, “Felt hopeless”, “Felt threatened, judged, or intimidated”, “Felt worthless or guilty”, and “Procrastinated”, for co-morbid and non-comorbid patients. The faceted plots show that mental symptoms are not static over time but rather volatile; there is autocorrelation between responses at two consecutive time points. Furthermore, the endorsement patterns also differ across patient diagnosis groups (COM vs. NON-COM). For example, co-morbid patients tend to have more severe responses to questions than the non-comorbid group, especially questions related to depression-like “Felt hopeless”, as well as those associated with anxiety-like “Procrastinated”; lastly, the temporal trend may vary between groups.

Fisher and Bosley (2020) suggests that the mood of mental disorder patients are likely to fall into discrete states. HDRBM is particularly useful for addressing this observation. The hidden layer with multiple discrete variables in HDRBM represents the unobserved mood states. In the EMA study, it can reveal differential moods across time and between patient groups. The model thus may identify distinct patterns of mood and behavior that may be difficult to discern using traditional latent variable methods such as factor analysis.

### 4.2 Analysis results

We compared the performances of the location-scale LMM and the proposed HDRBM on this EMA dataset. We fit an LMM model with one latent variable and random location intercepts, using all baseline measurements of observed variables as predictors. We allowed the variance (scale) of the random location intercepts to be different between diagnosis groups. Since the LMM model cannot handle discrete data, we used the original measures to fit the model. The final predictions were then obtained by discretizing the continuous predictions based on the same baseline tertiles used in our proposed method.

True observed measures (horizontal axis) and the predictions (vertical axis) from location-scale LMM are shown in Web Figure 3. Due to the shrinkage effect of mixed-effects models in general, location-scale LMM tends to shrink the values of its predictions towards population mean, such that the range of predictions is much smaller than the observed measures. In addition, since LMM assumes an ordinal relationship between the continuous latent variable and categorical outcomes, which is not supported by the observed data (e.g., orange points in Web Figure 3 did not conform with an ordinal relationship and were misclassified), the accuracy is much lower than our method which does not require the ordinal assumption. As shown in Table 1, our model shows significant improvement in both training and test accuracy over the LMM model. The test accuracy of HDRBM was computed from the average 5-fold cross-validation of the EMA data. The predictions were stable across folds, suggesting that it is robust and likely to be more generalizable to new samples. To further examine the quality of HDRBM as a generative model, we compare observed measures and data independently generated from fitted HDRBM in Figure 3. The results show that the distribution of all measures over time reconstructed from HDRBM closely resembles the original observed data, demonstrating the effectiveness of using HDRBM as a model for representing EMA data.

**Table 1:** Mean and standard deviation of the training and test accuracy of location-scale LMM and HDRBM on the real EMA data over 31 days (5-fold CV).

|  | LMM | HDRBM |
| --- | --- | --- |
| Training | 0.465 (0.019) | 0.563 (0.049) |
| Testing | 0.353 (0.089) | 0.536 (0.063) |

The bootstrap results shown in Web Table 6 indicate the presence of heterogeneity of dynamic structure between diagnosis groups in our EMA study, which cannot be captured by a simple RBM or RTRBM model. We illustrate the interpretability of our model results on the EMA study. Figure 4 shows the 95% bootstrap confidence intervals of the log odds ratios, which can be obtained from ***Ŵ*** with the reference level as “mild”. For any latent node *i* and visible node *j*, the log odds ratio between two different levels *k*_1_ and *k*_2_ *∈ {*1, 2, 3*}* can be expressed as the difference between corresponding weights, 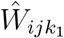 *−* 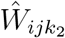 .

The subplots of the two hidden nodes (Figure 4) display questions related to negative mental symptoms arranged in increasing log odds ratios for the “severe” group from top to bottom. A higher odds ratio for a question indicates a stronger association between the activation of the hidden node and the deteriorating symptoms of that question. For instance, hidden node 1 is most strongly associated with the severe signals of negative symptoms of questions like “Felt down or depressed,” “Experienced loss of interest or pleasure,” and “Felt worthless or guilty,” while hidden node 2 is associated with negative symptoms of “Felt hopeless,” “Felt down or depressed,” and “Felt worthless or guilty.” One interesting observation is that the first hidden node expresses deteriorating symptoms of some reverse-coded questions, such as “Didn’t feel positive,” “Didn’t feel content,” and “Didn’t feel enthusiastic,” which are less strongly associated with the second hidden node. We can also examine weights directly to gain more insight into the associations between the latent and observed variables. In Web Figure 4, a node exhibiting a pattern of red (negative) to blue (positive) indicates that the activation of that node implies endorsing deteriorating symptoms for the questions with more saturated colors (which correspond to larger weights).

HDRBM also allows us to make inferences about the heterogeneous group dynamics of temporal dependencies between consecutive hidden states. Given the hidden node activation probabilities at time *t −* 1 and parameter estimates of **V**, we obtain

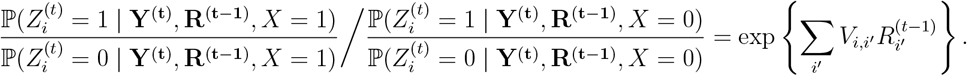

According to the above expression, the co-morbid group will have a higher odds of 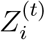 activation than the non-co-morbid group if and only if 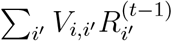 is greater than zero,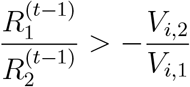, when there are two hidden nodes. Note that since 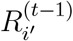 is the activation probability of hidden node *i*^*′*^ at the previous timepoint, it is always greater than or equal to zero. Consider our estimated (*V*_1,1_, *V*_1,2_) of (0.45, 0.14), the means in Web Table 6 of bootstrapped confidence intervals. These estimates suggest that the co-morbid group has higher odds of hidden node one activation at time *t* than the non-co-morbid group. Similarly, our estimated (*V*_2,1_, *V*_2,2_) are both positive (See Web Table 6), implying that the co-morbid group has a higher odds of hidden node two activations at time *t* as well and a higher temporal dependence over time. In summary, the results suggest that over time, the co-morbid group has higher odds of experiencing more severe symptoms and a higher autocorrelation than the depression-only or anxiety-only group, consistent with clinical observations.

Furthermore, hidden node activation probabilities can be obtained for each subject at every time point and be used to cluster patients. So with two hidden nodes at each time point, a patient can be clustered into one of four clusters. For each hidden node and every subject, we calculated the average hidden node activation probability over the entire month and obtained four clusters based on those probabilities, (0,0), (0,1), (1,0), and (1,1). See Web Table 7 for demographic and after treatment clinical characteristics by cluster and Web Figure 6 for mean trend and 95% quantiles of items over time by cluster. It is clear that for most items, cluster 1 (blue) has the lowest mean trend over time (RMSSD = 14.62), and cluster 4 (red) has the highest mean trend over time (RMSSD = 15.09). Cluster 1 consists of subjects whose hidden nodes are, on average, off for both hidden nodes over a month, and cluster 4 consists of subjects whose hidden nodes are, on average, on for both nodes over a month. Clusters 2 (RMSSD = 18.11) and 3 (RMSSD = 14.68) are groups whose subjects have one of two hidden nodes activated on average. Cluster 2 appears to be more variable over time.

Figure 5 presents four radar plots showing the average responses for each cluster across various emotional well-being categories: ‘Depression and Avoidance Behavior’, ‘Emotional Distress’, ‘Positive Emotions (reversed)’ and ‘Sleep and Physical Discomfort’. We observe similar findings to that of Web Figure 6 in that cluster 1 subjects show the least severe responses and cluster 4 subjects show the most severe responses. Cluster 2 subjects display more variability across items than other clusters, especially in the categories ‘Depression and Avoidance Behavior’ and ‘Sleep and Physical Discomfort’. Some items that display more severe responses in each of the emotional well-being categories are “Felt worthless or guilty” and “Had difficulty concentrating” (Depression and Avoidance Behavior), “Felt threatened, judged, or intimidated” (Emotional Distress), and “Dwelled on the past”, “Felt fatigued” and “Experienced muscle tension” (Sleepy and Physical Discomfort). These clusters also associate with differential treatment outcomes measured by HAMD change scores and HAMA change scores (Web Table 7).

**Figure 5:**
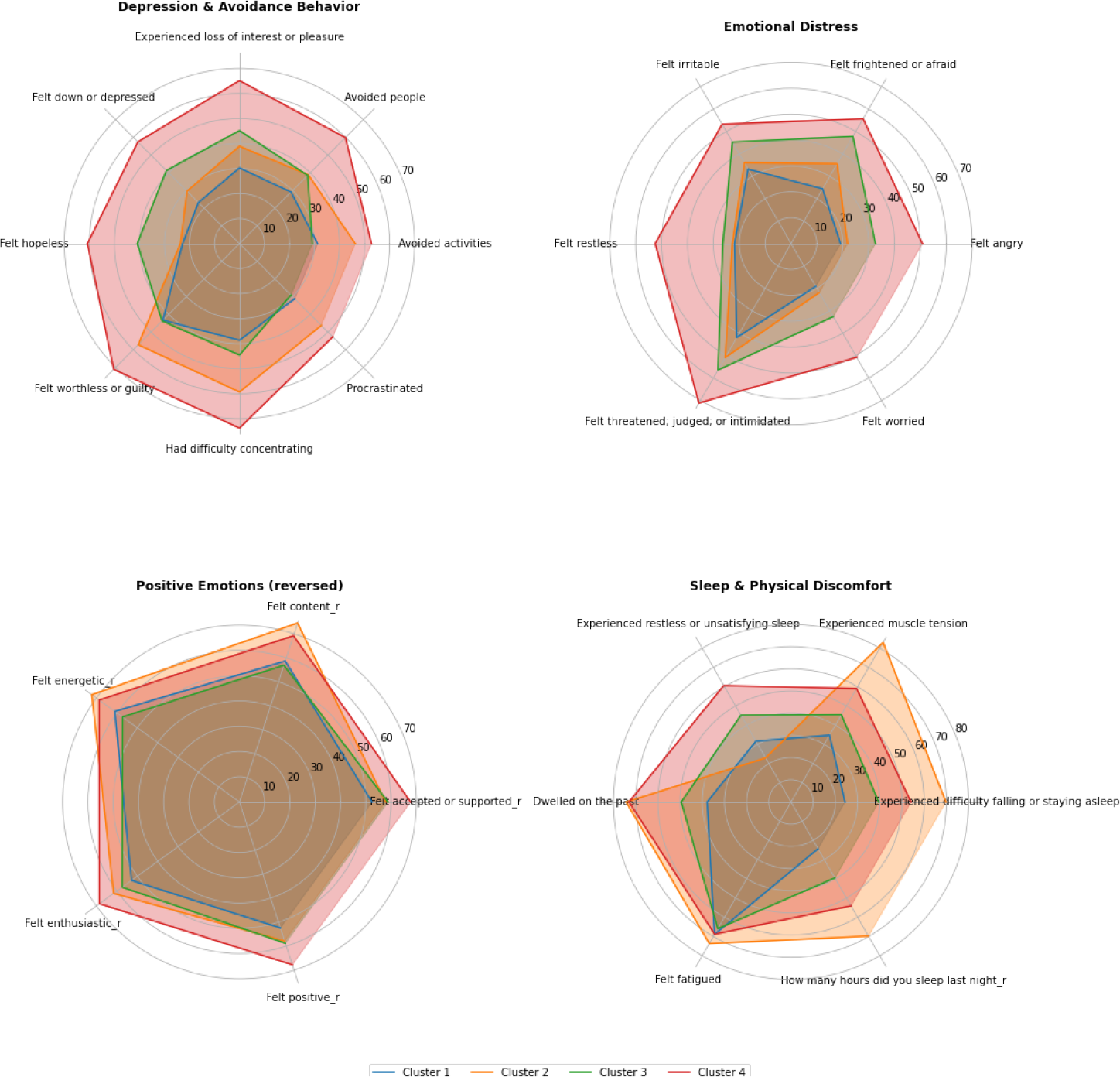
Radar plots showing the average responses for each cluster across various emotional well-being categories of the EMA study (Note: Four item categories are obtained from factor analysis).

## 5. Discussion

The combination of measurement theory, machine learning, and real-time data collection through EMA has the potential to improve the assessment of mental disorders and enable more personalized and effective care. In this work, we propose a temporal generative model, HDRBM, to learn the heterogeneity of group temporal dynamics for EMA data. Existing statistical methods for analyzing EMA data are variations of the mixed-effects model. However, when there are multiple discrete measures collected from time-intensive EMA studies on a heterogeneous population, a much more flexible modeling strategy is needed to fully address the complexity. By allowing temporal dynamics to depend on covariates, our HDRBM captures how they differ between groups with different symptom profiles. Furthermore, the interpretability of the model can provide information on the underlying drivers of mood experiences. Under the generative model framework, we can use HDRBM to impute missing data or simulate realistic synthetic EMA data for designing future studies, for example, clinical trials using EMA as outcome measures.

With increased layers and hidden nodes, HDRBM is expected to achieve universal approximation (Odense and Edwards, 2016). This extension to deep generative models is worth considering when the sample size is sufficient. Although the number of measures and time points in an EMA study may be large, the number of subjects is often small. A practical solution is to consider using data augmentation to improve parameter learning and prediction performance or leverage multiple studies. Our method can be easily applied to studies with multiple time-varying covariates and extended to multi-dimensional continuous measures by using Gaussian RBM (Cho et al., 2011) as building blocks. Furthermore, it is worth examining the theoretical identifiability of HDRBM along the lines of Cueto et al. (2010). Lastly, future work can also explore the use of HDRBM in other applications beyond EMA data, such as wearable devices, where time-varying covariates and heterogeneous temporal dynamics are commonly observed.

## Acknowledgements

This research was supported by U.S. NIH grants NS073671, GM124104, MH123487, and MH124106. We would like to thank authors in Fisher and Bosley (2020) for sharing their research data at the Open Science Framework (OSF). The EMA data in this work can be obtained at: https://osf.io/34xyh/.

